# A Floxed exon (Flexon) approach to Cre-mediated conditional gene expression

**DOI:** 10.1101/2021.07.13.452276

**Authors:** Justin M. Shaffer, Iva Greenwald

**Affiliations:** Dept. of Biological Sciences, Columbia University, New York, NY 10027

**Keywords:** conditional gene expression, lox-stop-lox, stop cassette, *C. elegans*

## Abstract

Conditional gene expression allows for genes to be manipulated and lineages to be marked during development. In the established “lox-stop-lox” approach, Cre-mediated tissue-specific gene expression is achieved by excising the stop cassette, a *lox*-flanked translational stop that is inserted into the 5′ untranslated region of a gene to halt its expression. Although lox-stop-lox has been successfully used in many experimental systems, the design of traditional stop cassettes also has common issues and limitations. Here, we describe the Floxed exon (Flexon), a stop cassette within an artificial exon that can be inserted flexibly into the coding region of any gene to cause premature termination of translation and nonsense-mediated decay of the mRNA. We demonstrate its efficacy in *C. elegans* by showing that, when promoters that cause weak and/or transient cell-specific expression are used to drive Cre in combination with a *gfp(flexon)* transgene, strong and sustained expression is obtained in specific lineages. We also describe several potential additional applications for using Flexon for developmental studies, including more precise control of gene expression using intersectional methods, tissue-specific protein degradation or RNAi, and generation of genetic mosaics. The Flexon approach should be feasible in any system where any site-specific recombination-based method may be applied.

**Summary statement:** The Floxed exon (Flexon), a stop cassette that can be inserted flexibly into the coding region of any gene, facilitates Cre-mediated conditional gene expression.

## INTRODUCTION

Conditional gene expression is an important tool for understanding how genes and proteins function in different tissues, lineages, and stages of development in complex organisms. Traditionally, control of gene expression is achieved by using regulatory regions derived from genes that are expressed in specific spatial and temporal patterns. However, the characterized regulatory regions do not always have the ideal expression pattern, and often trade-offs must be made between expression level and spatiotemporal specificity.

We have developed a method to address some of these limitations by devising a variation on the “lox-stop-lox”, or stop cassette, strategy that has been widely used in many systems, including *C. elegans* (van der Vaart et al., 2020). In the typical lox-stop-lox application, stop cassettes consist of one or more stop codons, each followed by a 3’ UTR. The full sequence is flanked by lox sites, and inserted into the 5’ UTR of a gene of interest to block expression of the gene (Fig 1A). When Cre recombinase is provided through a separate driver, the cassette is excised and the gene of interest can be expressed (Lakso et al., 1992). Stop cassettes are used in transgenes to generate spatiotemporal specificity via tissue-specific Cre drivers, and high expression levels by placing the cassette between the coding region and a strong promoter. For example, conditional control of transgene expression has been accomplished in mice by inserting transgenes with stop cassettes into the Rosa26 locus, a neutral locus with the potential for mid-level expression in multiple tissues after Cre-mediated excision (Friedrich and Soriano, 1991; Hitoshi et al., 1991; Nagy, 2000). The Flp-FRT system of site-specific recombination has also been used to excise stop cassettes (Dymecki, 1996).

**Figure 1.**
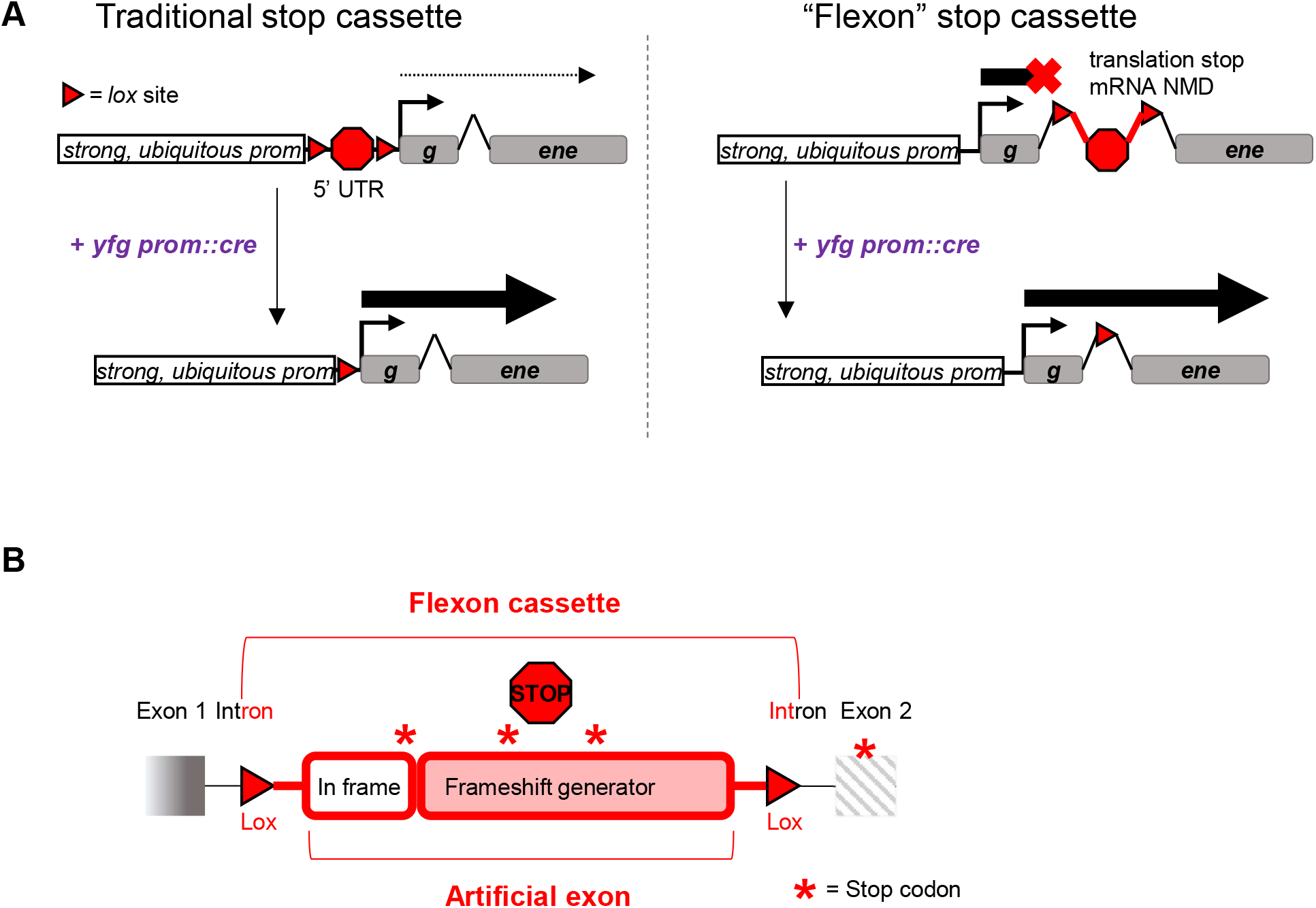
Design and use of a <u>Fl</u>oxed <u>ex</u>on (*flexon*) cassette. (A) A traditional stop cassette consists of one to three stop codon and 3’ UTR sequences, with the entirety of the sequence flanked by *lox* sites. The *flexon* stop cassette consists of an artificial exon that contains an in-frame stop codon followed by a frameshift generator that will create stop codons in all potential reading frames, flanked by introns bounded by *lox* sites. In the application described herein, the stop cassette is incorporated into a transgene to block expression from a strong, ubiquitous promoter. Tissue-specific expression of Cre recombinase, driven by the promoter of “*your favorite gene*” (*yfg*), results in tissue-specific restoration of gene expression. (B) The artificial exon of the *flexon* cassette used to construct the *gfp(flexon)* gene tested in Figs. 2 and 3 has an in-frame, three codon leader sequence that ends with an in-frame stop codon. The following sequence contains additional stop codons in all reading frames and generates downstream frameshift mutations in all frames of the coding region. The *flexon* used in this study is bounded by *lox2272* sites.

Traditional stop cassettes have several common issues and limitations. Ectopic expression of the gene can occur because the start site and entire open reading frame are intact. This leaky expression is compounded when using cassettes with stop sequence repeats because low, basal levels of Cre expression can result in partial recombination and weakened gene repression (Bapst et al., 2020). While leaky expression is the chief concern for stop cassette usage in transgenes, further limitations apply when using a stop cassette in an endogenous context. A situation common to many organisms is that different isoforms of a gene have varying start sites, potentially complicating the application of traditional lox-stop-lox to create likely null alleles. The 5’ UTR may also contain uncharacterized but important regulatory sequences; in *C. elegans*, these could also include trans-splicing signals upstream of conventional genes and internal to operons (reviewed in Arribere et al., 2020). Placing a stop cassette into these sequences could alter expression of the downstream gene; even after the cassette is excised, the lox scar could disrupt important regulatory sequences.

Here, we outline a strategy for a stop cassette that cuts down on leaky expression and increases insertion site flexibility for both transgenes and endogenous genes, while retaining spatiotemporal control of expression. We call this strategy “Flexon,” for “Floxed exon,” in which we engineer a stop cassette in the form of an artificial exon that can prevent protein translation by causing premature termination in all three reading frames. The Flexon cassette can be inserted into an intron or exon, providing different options for its placement within the coding region of a transgene or endogenous locus. We demonstrate the efficacy of Flexon as a stop cassette for tissue-specific gene expression in specific lineages, and describe other potential applications of Flexon for genetic analysis.

## RESULTS

### Design considerations

#### General design of the Floxed exon (Flexon) cassette

As with traditional stop cassettes, the principle behind the Flexon cassette is that it should prevent target gene expression in the absence of Cre-mediated excision, so that tissue-specific expression of Cre will excise the cassette and restore the open reading frame in the tissue of interest. In addition, the Flexon cassette is designed to (i) be versatile regarding where it can be inserted in an open reading frame, and (ii) prevent ectopic gene expression due to spurious initiation downstream of the cassette, which is an issue with traditional stop cassettes.

The Flexon cassette contains a stop codon in an artificial exon flanked by artificial introns for flexible placement within a coding region (Fig. 1A). Each flanking intron contains a single lox site, and both lox sites are oriented in the same direction to allow for excision of the stop sequence in the exon (Dickinson et al., 2015). Due to the flanking introns, the stop sequence is spliced into the transcript if the cassette is inserted into an exon or an intron.

The artificial exon contains redundant mechanisms to halt translation of the protein: a short nucleotide sequence with an in-frame stop codon relative to the gene of interest, followed by a longer nucleotide sequence that results in a frameshift and stop codons in the other reading frames (Fig. 1B). If the initial premature stop is read through, the subsequent frameshift mutation will also trigger nonsense-mediated decay or create a non-functional protein. When Cre-mediated recombination occurs, the exon with the stop sequence is removed from the DNA sequence, leaving a single artificial intron with a *lox* scar in the middle and restoring the reading frame to encode for a functional protein (Fig. 1A).

#### Design of a “*gfp(flexon)”* cassette

To test the Flexon approach, we incorporated a Flexon cassette into GFP for Cre-dependent expression in *C. elegans*, using the following specific sequences. We present general design considerations that could be applied to other Flexon cassettes in *C. elegans* or other experimental systems in the Discussion.

1. In-frame exon sequence: the exon contains a three-codon leader to a stop codon that is in frame with the coding region for GFP (Fig 1B).
2. Frameshift-generating exon sequence: because traditional stop cassettes use single or repetitive 3’UTR sequences (Lakso et al., 1992; Madisen et al., 2010; Maxwell et al., 1989; van der Vaart et al., 2020), we used 62 bp of sequence from the neutral 3’ UTR from *tbb-*2 (Merritt et al., 2008) to cause a frameshift of the downstream coding region (Fig 1B). The total length of the exon sequence is the same as the 2nd exon in a widely-used, artificial intron-containing form of *gfp* (Dickinson et al., 2015) to ensure it is of sufficient length to be spliced into the mRNA. The length of the exon was kept as small as possible to ensure efficient Cre-mediated excision, and the short length also facilitates cloning and homologous repair for insertion into an endogenous gene.
3. Intron sequences: the artificial introns were derived from a self-excising drug selection cassette (Dickinson et al., 2015). Artificial introns of similar sequences (Fire Vector Kit, 1995) are commonly used for transgenes in *C. elegans* because they demonstrate efficient splicing, due to the short sequence length and strong splice acceptor and donor sequences.
4. *lox* site selection: we used the *lox* variant *lox2272* to avoid recombination with *loxP* sites in existing transgenes or engineered loci we routinely use in our work.
5. Cassette insertion: We replaced the first intron in a codon optimized GFP sequence (see Materials and Methods) to create a “*gfp(flexon)*” sequence.

### Testing the Flexon approach using a *gfp(flexon)*transgene for strong, Cre-mediated tissue-specific expression of GFP

We used the strong ubiquitous promoter *rps-27p* (Giordano-Santini et al., 2010) to drive expression of *gfp(flexon)* in all somatic cells. The resulting *rps-27p*::*gfp(flexon)* transgenes produce no or minimal visible GFP fluorescence on their own (Fig. S1), indicating little to no bypass expression of the stop cassette. We then combined *rps-27p::gfp(flexon)* with two different Cre drivers that were made using tissue-specific promoters that result in weak and/or transient expression. The results were remarkable: the expected lineages were permanently and brightly marked.

#### *Test of* gfp(flexon) *in the somatic gonad*

A *C. elegans* L1 larvae hatches with two somatic gonad precursor cells, Z1 and Z4, which generate all of the structures of the somatic gonad during postembryonic development (Fig. 2A). During the first phase of gonadogenesis, Z1 and Z4 give rise to twelve cells that form the somatic gonad primordium in the L2 stage, and the somatic gonad blast cells divide in the L3 stage and give rise to many additional cells (Kimble and Hirsh, 1979).

**Figure 2.**
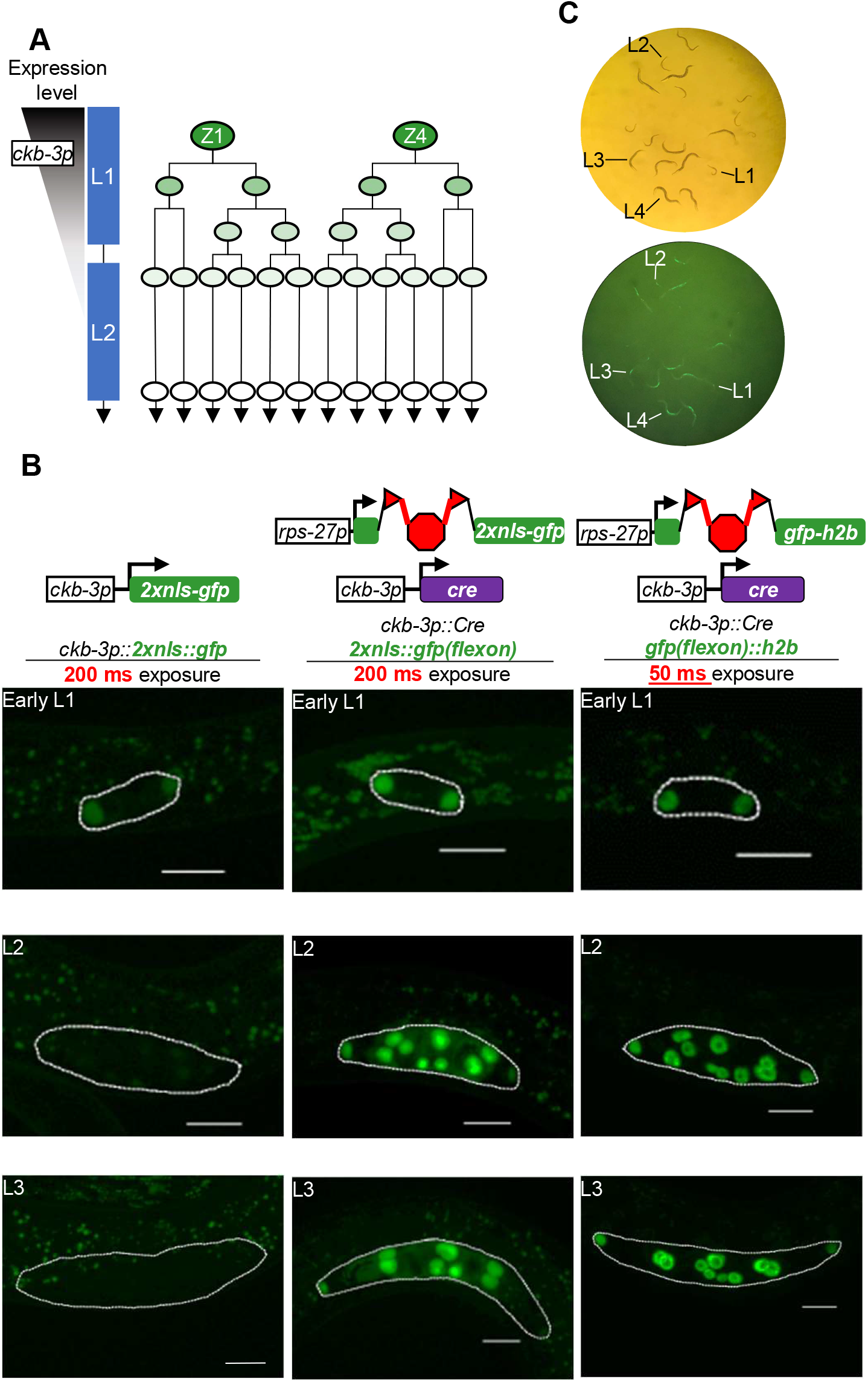
Flexon enables strong, persistent expression of GFP in all cells of the somatic gonad lineage. (A) *ckb-3p* is a highly-specific promoter for the somatic gonad precursor cells Z1 and Z4 (Kroetz and Zarkower 2015). (A, B) When used to express 2xnls::GFP (*arTi433[ckb-3p::2xnls::gfp])*, GFP is visible in the Z1 and Z4 nuclei, but decreases as the lineage proceeds and is undetectable by the time the somatic gonad primordium forms in the late L2 stage. (B) When *flexon* excisions were mediated by a *ckb-3p::cre* driver [*arTi237*; see Materials and Methods], GFP fluorescence intensity from *arTi435[rps-27p::2xnls::gfp(flexon)]* appears brighter in Z1 and Z4, and persists strongly throughout the gonad lineages, as seen in photomicrographs (middle column) taken at the same exposure time and imaging parameters that resulted in low or undetectable expression of *2xnls::gfp*. Stabilization of GFP using a histone tag (*arTi361[rps-27p::gfp(flexon)::h2b],* right column) results in bright expression visible at a lower exposure time than 2XNLS::GFP. All scale bars are 10 μm. (C) GFP::histone is visible on the dissecting scope at all larval stages. Magnification is 50x

The 5’ regulatory region for the *ckb-3* gene, denoted *ckb-3p*, drives expression in the somatic gonad precursor cells Z1 and Z4 in embryos and L1 larvae, but expression diminishes as the lineage progresses (Kroetz and Zarkower, 2015). When *ckb-3p* is used to drive 2xNLS::GFP, nuclear GFP is readily visualized in Z1 and Z4 in the L1 stage, but progressively dims and is essentially undetectable by the time the somatic gonad primordium has formed in the L2 stage (Fig. 2A, B). When combined with a *ckb-3p::Cre* driver (see Materials and Methods) that expresses a form of Cre recombinase optimized for *C. elegans* (Ruijtenberg and van den Heuvel, 2015), *rps-27p::2xnls::gfp(flexon)* produces sustained strong and specific expression of 2xNLS::GFP in the somatic gonad throughout development (Fig. 2B). In addition, Z1 and Z4 appeared brighter in the L1 stage when compared to *ckb-3p::2xnls::gfp* (Fig. 2B). Furthermore, when we used a histone tag to stabilize GFP as well as for nuclear localization (r*ps-27p::gfp(flexon)::h2b*), fluorescence intensity was sufficient for visualization at lower exposure, and even using a dissecting microscope (Fig. 2B, C).

#### *Test of* gfp(flexon) *in the vulval precursor cells*

The vulval precursor cells (VPCs) are six polarized epithelial cells, named P3.p-P8.p, that are born in the L1 stage and remain quiescent until the L3 stage. At that time, P5.p, P6.p, and P7.p are induced by EGFR and LIN-12/Notch signaling to adopt vulval fates; the descendants of these cells form the vulval primordium in the L4 stage (Gauthier and Rocheleau, 2017; Shin and Reiner, 2018; Sternberg, 2005). The other VPCs, P3.p, P4.p and P8.p, do not receive spatial patterning signals and divide once to generate daughter cells that fuse with the major hypodermal syncytium (P3.p sometimes fuses directly, without dividing). A composite regulatory region made from the 5’ flanking region and introns of the *lin-31* gene, called *lin-31p*, drives fluorescent reporter expression in all six VPCs and displays a dynamic pattern when the vulval fates are induced and executed (de la Cova and Greenwald, 2012; Luo et al., 2020; Tan et al., 1998; Fig. 3A). This dynamic pattern is evident using a single-copy insertion *lin-31p::2xnls::gfp* transgene. GFP fluorescence is visible after the cells are born in the late L1 stage and remains approximately uniform in all VPCs during the L2 stage. At the time VPCs commit to their fates in the L3 stage, fluorescence begins to decrease visibly in P5.p, P6.p, and P7.p and progressively dims and becomes undetectable as their lineages progress (Luo et al., 2020; Fig. 3B). Fluorescence in P3.p, P4.p and P8.p, which do not adopt vulval fates, remains stronger than in the other VPCs at the beginning of the L3 stage but becomes undetectable after they divide in the L3 stage. By contrast, when *rps-27*::*2xnls::gfp(flexon)* was combined with a *lin-31p::Cre* driver, GFP was strongly, uniformly, and continuously expressed in the VPCs and their descendants (Fig. 3B,C). As in the somatic gonad, the use of a histone tag to stabilize GFP results in fluorescence visible using a dissecting microscope (not shown).

**Figure 3.**
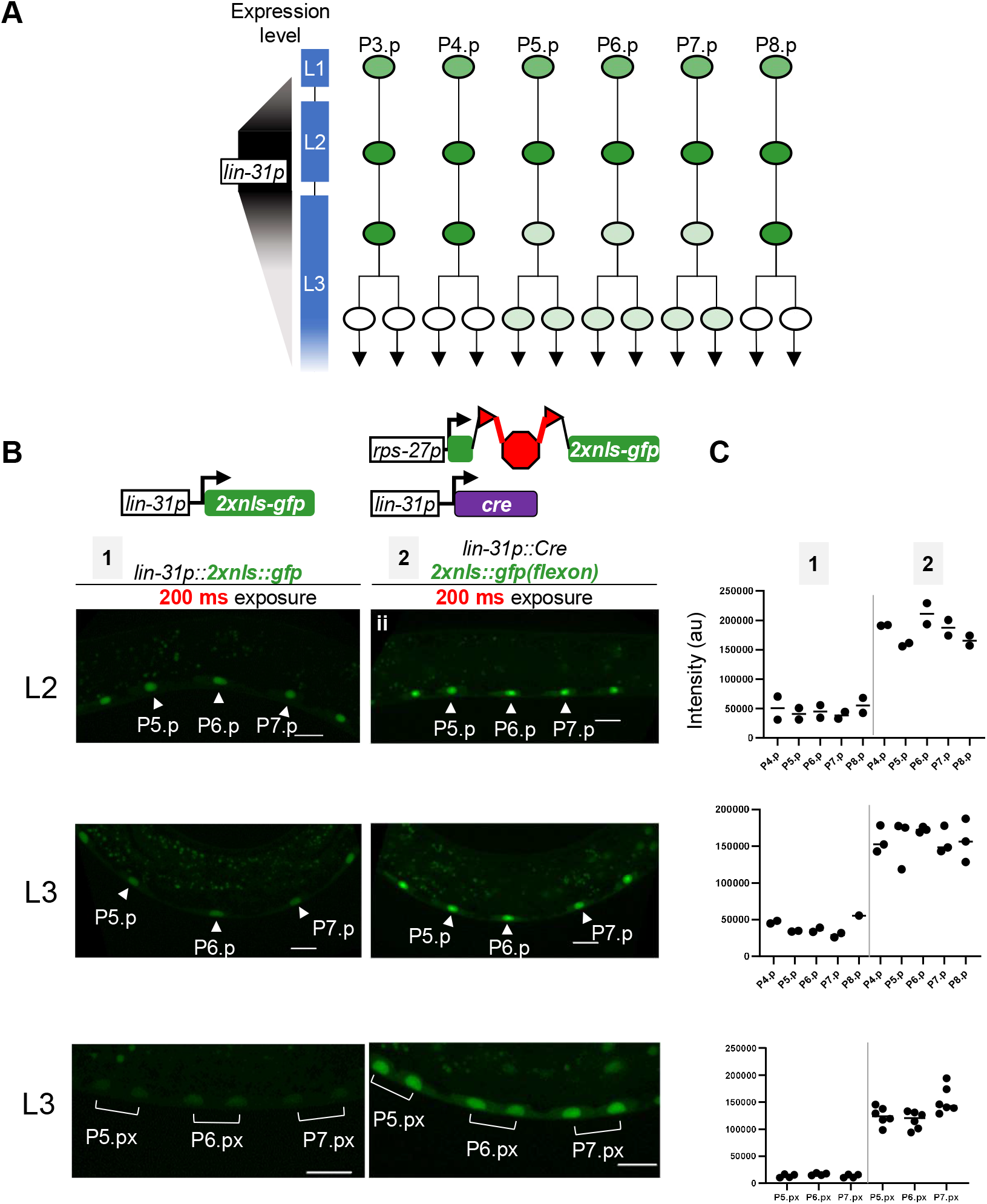
Flexon enables strong, persistent expression of GFP in the vulval precursor cell (VPC) lineages. (A) The *lin-31p* regulatory sequence drives expression in all VPCs in late L1 and L2, but becomes visibly non-uniform during vulval fate patterning in L3 and wanes as the lineages progress (de la Cova and Greenwald 2012; Luo *et al.* 2020; Tan *et al.* 1998). (B)Photomicrographs of *arTi434[lin-31p::2xnls::gfp]* (left column, gernotype 1), showing dynamic expression pattern of *lin-31p* schematized in A. By contrast, GFP fluorescence intensity from *arTi435[rps-27p::2xnls::gfp(flexon)]* (right column, genotype 2) is higher starting early in development, and persists strongly throughout the VPC lineages. These *flexon* excisions were mediated by the VPC Cre driver *arTi381[lin-31p::cre]*. All scale bars are 10 μm. (C) Quantitation of several individuals of genotypes 1 and 2 shown in (B).

## DISCUSSION

We demonstrated how the Flexon approach may be used for strong, tissue- or lineage-specific expression of GFP using weaker, more transiently expressed Cre drivers, thereby providing a prototype for strong, tissue-specific expression of any desired protein. In this Discussion, we first generalize the design considerations for other Flexon cassettes and then provide examples of how Flexon could be adapted to improve the efficiency of commonly used genetic tools and for other genetic approaches in *C. elegans* and other experimental systems.

### Design considerations for other Flexon cassettes

In principle, any protein-coding gene can be converted into a Flexon gene. Both the exon and *lox* sites are modifiable to suit different target genes. In Fig. 4A, we show several adjustments that may be made to the Flexon cassette, and highlight here considerations that should be borne in mind when customizing the Flexon. In addition to cassette adjustments, when a Flexon is used in a transgene, different ubiquitous or more restricted promoters may be used to tune the level or tissue in which the Flexon construct is expressed after excision of the cassette (Fig. 4B).

1. The beginning of the artificial exon should be adjusted to ensure that the frame is maintained until the in-frame premature stop codon.
2. The frameshift-generating sequence should be tested by conceptual translation to ensure that it causes a frameshift downstream, and may be modified to include additional out-of-frame stop codons within the exon if necessary.
3. The specific pair of *lox* sequences selected to flank the exon can be chosen to allow for the possibility to excise multiple Flexon cassettes simultaneously, or to guard against causing intergenic DNA recombination in the presence of other genes with *lox* scars from prior manipulations (such as the *loxP* scar left from the self-excising cassette method of genome engineering in *C. elegans*; Dickinson et al., 2015). Alternatively, FRT or other site-specific recombinase sequences can be used in place of *lox* sequences for spatiotemporal control of recombination and/or for differential control when combining different Flexon transgenes.
4. Insertion of Flexon into an exon can be achieved by appending a splice donor site upstream of the first lox sequence, and branch and splice acceptor sequences downstream of the second lox site. Sequence may also be added between the appended sequences and the lox sites to create introns of a certain size for high splicing efficiency in the organism of interest.

**Figure 4.**
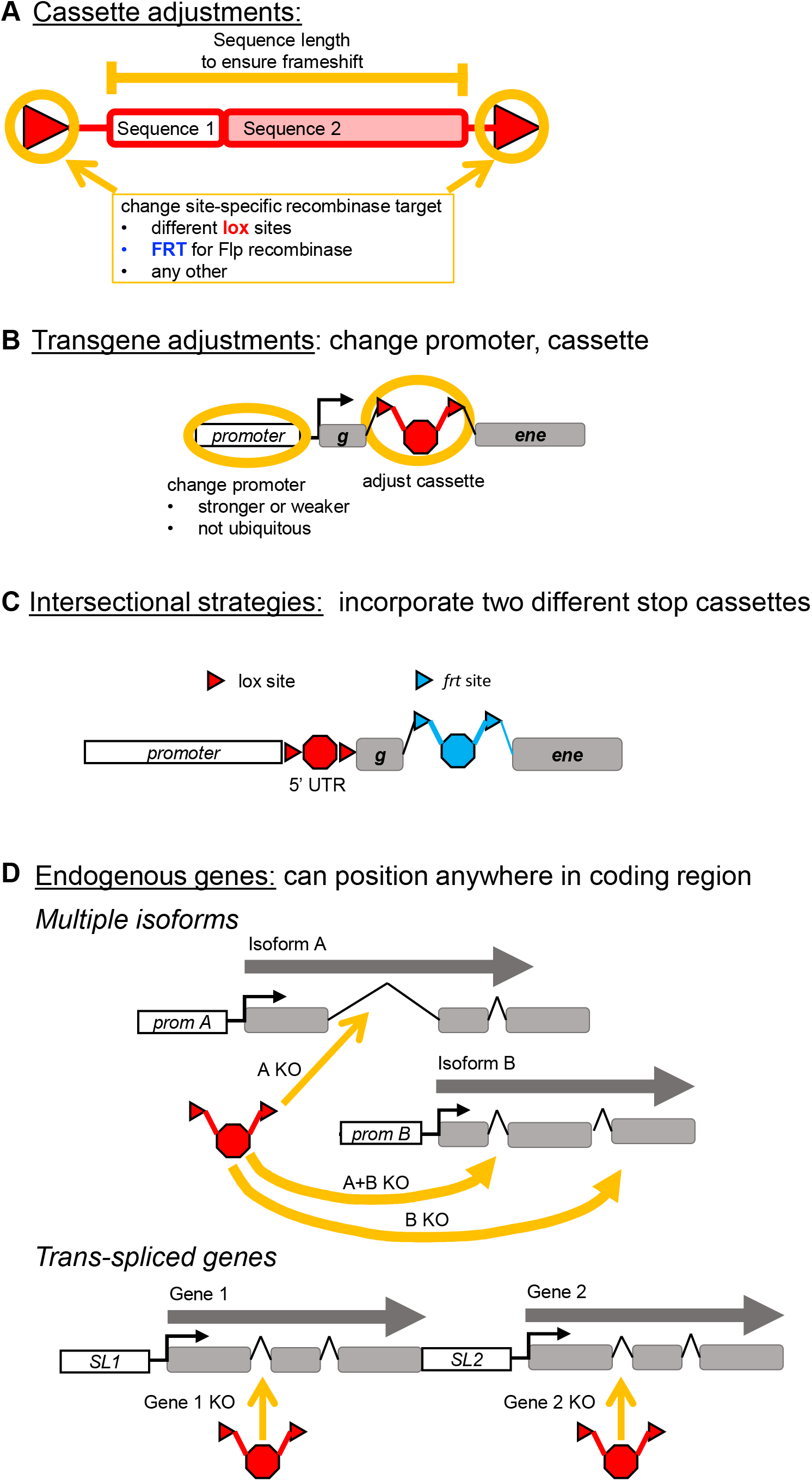
The Flexon system is tunable and may be incorporated into transgenes or endogenous loci to enable or improve genetic tools and approaches as well as to mark lineages. Adjustments to the design for specific purposes (A, B) and some potential applications (C, D) are diagrammed here and described in the text.

### Potential applications facilitated by incorporating Flexon into endogenous genes or transgenes

Flexon may be incorporated into many different established methods for manipulating gene activity, for studies of development or any other purposes. We give some examples here, emphasizing *C. elegans* but applicable to other experimental systems.

#### Intersectional approaches to conditional gene expression

One way to achieve specific gene expression is to use two different tissue specific promoters that have overlapping sites of expression to drive different recombinases, e.g. combining the Cre and Flp recombinases (Anastassiadis et al., 2010; Awatramani et al., 2003; Liu et al., 2020; Fig. 4C). The flexible placement of the *flexon* may facilitate the use of such intersectional strategies (Fig. 4D). Combinatorial recombinase approaches have been used for marking specific cell types and lineages in mice (reviewed in Dymecki et al., 2010); in *C. elegans*, these approaches may be used to generate GFP lineage markers that require lower intensity and exposure to visualize, avoiding damage from blue-light overexposure (Edwards et al., 2008; Ward et al., 2008).

#### Increasing the level of limiting effectors of tissue-specific protein degradation or RNAi

Targeted protein degradation using natural degron/ubiquitin ligase pairs, such as the TIR1/AID system [applied to *C. elegans* by (Zhang et al., 2015)] and the ZIF-1/ZF1 system (Armenti et al., 2014), or converting GFP itself into a degron by replacing the interaction domain of an E3 ubiquitin ligase with an anti-GFP nanobody (Wang et al., 2017), are powerful approaches to studying gene function. However, the amount of the ubiquitin ligase expressed has been reported to be limiting for degradation (Aghayeva et al., 2021; Ashley et al., 2021; Dos Santos Maraschin et al., 2009), so we anticipate that the Flexon approach will facilitate such manipulations by enabling stronger tissue-specific expression of the ubiquitin ligase. Furthermore, by using different *lox* variants, Cre can be used to delete a floxed, degron-tagged gene while simultaneously using a *ubiquitin ligase(flexon)* to quickly eliminate perduring protein, a strategy that has been demonstrated to be effective when low residual protein levels obscure the null phenotype (van der Vaart et al., 2020).

Similarly, the Flexon approach may allow for improved tissue-specific RNAi using the established approach of tissue-specific rescue of a mutation defective in RNAi such as *rde-1* (Qadota et al., 2007; Watts et al., 2020) or *sid-1* (Kage-Nakadai et al., 2014; Winston et al., 2002). Alternatively, a universal tissue-specific RNAi strain could in principle be made by inserting a Flexon cassette into the endogenous *rde-1* or *sid-1* gene, restoring physiological levels of expression of the gene product to specific cells after Cre excision.

#### Engineering genomic loci

For engineering genomic loci, a *flexon* offers several potential advantages over conventional lox-stop-lox cassettes. First, the Flexon approach facilitates engineering of alternatively spliced genes, as the cassette may be placed so as to abrogate expression of single or multiple isoforms (Fig. 4D, top). Second, in many cases introns are better defined than 5′ UTRs, especially in *C. elegans* where about 84% of genes are trans-spliced by the addition of a splice-leader sequence to the 5′ end of pre-mRNAs (Tourasse et al., 2017), and about 15% of genes are expressed from operons, in which a polycistronic pre-mRNA is processed into separate transcripts by trans-splicing of the downstream gene (Allen et al., 2011), in which the trans-splice acceptor for the downstream gene is embedded in a relatively short sequence that also contains a 3′ UTR for the upstream gene. Thus, a *flexon* can be inserted into the coding region of a trans-spliced gene, or into either gene of an operon, without potentially compromising regulatory sequences (Fig. 4D, bottom).

Finally, we note that manipulating endogenous gene expression using the Flexon approach offers an alternative to transgene-based methods for assessing tissue-specific rescue and creating genetic mosaics that may be advantageous in certain situations. For example, transgenes used for conventional tissue-specific rescue may not be expressed at endogenous levels, and when genes have multiple isoforms, the isoform selected may influence the results. By contrast, insertion of a Flexon cassette into an intron of an endogenous gene will allow for all isoforms that share that intron to be expressed under control of its natural regulatory elements after excision using a tissue-specific Cre driver (Fig. 4D, top).

### Potential Limitations

Although the Flexon approach addresses some of the issues that may arise with traditional lox-stop-lox, it shares other limitations inherent to any stop cassette method. One is that site-specific recombination is irreversible and therefore cannot be used for dynamic control of gene expression by itself, although it may facilitate the implementation of dynamic methods such as AID as described above. Another limitation is that transient or variable low-level expression that is not normally evident when the promoters are identified using fluorescent reporter genes may be more apparent when they are used to drive Cre (Tenen and Greenwald, 2019), so tissue specific promoters used to drive site-specific recombinases may not be as specific as desired. Finally, expression in different cells of a tissue may be uneven shortly after recombination events due to asynchronous cassette excision or when occasionally excision does not occur on both chromosomes in a cell. However, recombination tends to be very efficient given the small size of the *flexon*, so for many applications, this limitation may not be an issue.

## MATERIALS AND METHODS

### *C. elegans* strains

*C. elegans* was grown on 6 cm NGM plates seeded with *E. coli* OP50 and maintained at 20°C. Strain N2 (wild-type; Brenner, 1974) was previously described, and GS8795 *arTi237 [ckb-3p::Cre(opti)::tbb-2 3’ UTR] X* is a single-copy insertion transgene made as described in (Tenen and Greenwald, 2019), and mapped as part of this study. Cre(opti) refers to the codon-optimized Cre recombinase described in Ruijtenberg and van den Heuvel (2015).

The following strains were generated in this study. Strains containing two transgenes were constructed using conventional crosses, and confirmed using PCR and microscopy.

GS9443 *arTi381 [lin-31p::Cre(opti)::tbb-2 3’ UTR] V*
GS9407 *arTi361 [pHK001(rps-27p::gfp(flexon)::his-58::unc-54 3’ UTR)] I*
GS9684 *arTi433 [pJS110(ckb-3p::2xnls::gfp::unc-54 3’ UTR)] V*
GS9690 *arTi435 [pJS145(rps-27p::2xnls::gfp(flexon)::unc-54 3’ UTR)] I*
GS9686 *arTi434 [pJS146(lin-31p::2xnls::gfp::unc-54 3’ UTR)] IV*
GS9401 *arTi361 I; arTi237 X*
GS9532 *arTi361 I; arTi381 X*
GS9691 *arTi435 I; arTi381 V*
GS9692 *arTi435 I; arTi237 X*

### Generation of single-copy insertion transgenes

The plasmids pHK001 [*rps-27p::gfp(flexon)::his-58::unc-54 3’ UTR*], pJS110 [*ckb-3p::2xnls::gfp::unc-54 3’ UTR*], pJS145 [*rps-27p::2xnls::gfp(flexon)::unc-54 3’ UTR*], and pJS146 [*lin-31p::2xnls::gfp::unc-54 3’ UTR*] were made in the miniMos vector backbone pCFJ910 (Frøkjær-Jensen et al., 2014) using Gibson Assembly (NEB Inc., MA) and confirmed by sequencing. pJS110 and pJS146 contain a *C. elegans* codon-optimized GFP sequence tagged with an N-terminal SV40 and C-terminal *egl-13* nuclear localization sequences (*nls*), regulated by the neutral 3’ UTR from *unc-54* (Hunt-Newbury et al., 2007). Plasmids were injected into N2 hermaphrodites. Random, single-copy insertions were obtained and mapped using the standard protocol (Frøkjær-Jensen et al., 2014).

### Microscopy

*C. elegans* larvae were imaged on a Zeiss AxioObserver Z1 inverted microscope with a 63x, 1.4NA oil immersion objective equipped with a spinning disk, CSU-X1A, a laser bench rack, and a Photometrics Evolve EMCCD camera. For GFP fluorescent imaging, a 488nm, 100mW laser was used for excitation. Larvae were mounted onto 3% agar pads, and immobilized with 10 mM levamisole.

Z stacks were collected from GS9684, GS9686, GS9691, and GS9692, and GS9401 larvae (Figures 2 and 3) with slices at 500 nm intervals and was imaged for GFP fluorescence with the following parameters: 10% laser power, 200 ms exposure (or 50 ms exposure for GS9401), and 400 EM gain. For strains GS9684, GS9692, and GS9401, the number of slices used per larva varied with the size of the gonad to fully capture the nuclei of every somatic gonad cell present. The stage of the animal was determined by the number of somatic gonad cells and somatic gonad morphology. For GS9686 and GS9691, each stack contained 26 slices, which was sufficient to image the full volume of the nuclei of every VPC or VPC descendent. The stage of the animal was determined by the number of VPCs and somatic gonad morphology.

For Figure S1, Z stacks were collected from GS9401 and GS9407 with slices at 500 nm intervals and imaged for GFP fluorescence with the following parameters: 25% laser power, 500 EM gain, and the exposure time denoted in the figure. The number of slices used per larva varied with the size of the gonad to fully capture the nuclei of every somatic gonad cell present. The stage of the animal was determined by the number of somatic gonad cells and somatic gonad morphology.

Images in Figure 2C were taken on a phone camera through the eyepiece of a Zeiss Discovery V.12 SteREO dissecting microscope, GFP470 filter, Schott ACE 1 fiber optic light source, and X-cite series 120Q fluorescent lamp illuminator. Animals to be imaged were placed on 60 mm plates filled with 1.75% agarose, and then immobilized using 10 mM levamisole.

### Image Quantification

Fluorescence intensity was quantified using FIJI (Linkert et al., 2010; Schindelin et al., 2012). Z stacks were sum projected for all slices containing cells of interest. Nuclei were manually segmented, and the mean green fluorescent background from a random region outside of the animal was subtracted from the mean green fluorescent intensity of each nucleus to correct for background.

## ACKNOWLEDGEMENTS

We thank Claudia Tenen for generating *arTi237*, Katherine Luo for generating *arTi381*, and Jee Hun (Henry) Kim for help in generating *arTi361*. We also thank Michael Shen, Michelle Attner, and Catherine O’Keeffe for valuable comments on the manuscript. This work was supported by R35GM131746 from the National Institute of General Medical Sciences (to I.G.), F31EY030331 from the National Eye Institute (to J.M.S.). J.M.S. was also supported by training grant T32GM008798. Some strains were provided by the CGC, which is funded by NIH Office of Research Infrastructure Programs (P40 OD010440).

## References

Aghayeva, U., Bhattacharya, A., Sural, S., Jaeger, E., Churgin, M., Fang-Yen, C., Hobert, O., 2021. DAF-16/FoxO and DAF-12/VDR control cellular plasticity both cell-autonomously and via interorgan signaling. PLOS Biology 19, e3001204.

Allen, M.A., Hillier, L.W., Waterston, R.H., Blumenthal, T., 2011. A global analysis of C. elegans trans-splicing. Genome Research 21, 255–264.

Anastassiadis, K., Glaser, S., Kranz, A., Bernhardt, K., Stewart, A.F., 2010. Chapter Seven - A Practical Summary of Site-Specific Recombination, Conditional Mutagenesis, and Tamoxifen Induction of CreERT2, in: Wassarman, P.M., Soriano, P.M. (Eds.), Methods in Enzymology. Academic Press, pp. 109–123.

Armenti, S.T., Lohmer, L.L., Sherwood, D.R., Nance, J., 2014. Repurposing an endogenous degradation system for rapid and targeted depletion of *C. elegans* proteins. Development (Cambridge, England) 141, 4640–4647.

Arribere, J.A., Kuroyanagi, H., Hundley, H.A., 2020. mRNA Editing, Processing and Quality Control in *Caenorhabditis elegans*. Genetics 215, 531–568.

Ashley, G.E., Duong, T., Levenson, M.T., Martinez, M.A.Q., Johnson, L.C., Hibshman, J.D., Saeger, H.N., Palmisano, N.J., Doonan, R., Martinez-Mendez, R., Davidson, B.R., Zhang, W., Ragle, J.M., Medwig-Kinney, T.N., Sirota, S.S., Goldstein, B., Matus, D.Q., Dickinson, D.J., Reiner, D.J., Ward, J.D., 2021. An expanded auxin-inducible degron toolkit for Caenorhabditis elegans. Genetics 217.

Awatramani, R., Soriano, P., Rodriguez, C., Mai, J.J., Dymecki, S.M., 2003. Cryptic boundaries in roof plate and choroid plexus identified by intersectional gene activation. Nature Genetics 35, 70–75.

Bapst, A.M., Dahl, S.L., Knöpfel, T., Wenger, R.H., 2020. Cre-mediated, loxP independent sequential recombination of a tripartite transcriptional stop cassette allows for partial read-through transcription. Biochimica et Biophysica Acta (BBA) - Gene Regulatory Mechanisms, 1863, 194568.

de la Cova, C., Greenwald, I., 2012. SEL-10/Fbw7-dependent negative feedback regulation of LIN-45/Braf signaling in C. elegans via a conserved phosphodegron. Genes & development 26, 2524–2535.

Dickinson, D.J., Pani, A.M., Heppert, J.K., Higgins, C.D., Goldstein, B., 2015. Streamlined Genome Engineering with a Self-Excising Drug Selection Cassette. Genetics 200, 1035–1049.

Dos Santos Maraschin, F., Memelink, J., Offringa, R., 2009. Auxin-induced, SCFTIR1-mediated poly-ubiquitination marks AUX/IAA proteins for degradation. The Plant Journal 59, 100–109.

Dymecki, S.M., 1996. Flp recombinase promotes site-specific DNA recombination in embryonic stem cells and transgenic mice. Proceedings of the National Academy of Sciences 93, 6191–6196.

Dymecki, S.M., Ray, R.S., Kim, J.C., 2010. Chapter Eleven - Mapping Cell Fate and Function Using Recombinase-Based Intersectional Strategies, in: Wassarman, P.M., Soriano, P.M. (Eds.), Methods in Enzymology. Academic Press, pp. 183–213.

Edwards, S.L., Charlie, N.K., Milfort, M.C., Brown, B.S., Gravlin, C.N., Knecht, J.E., Miller, K.G., 2008. A Novel Molecular Solution for Ultraviolet Light Detection in Caenorhabditis elegans. PLOS Biology 6, e198.

Friedrich, G., Soriano, P., 1991. Promoter traps in embryonic stem cells: a genetic screen to identify and mutate developmental genes in mice. Genes & development 5, 1513–1523.

Frøkjær-Jensen, C., Davis, M.W., Sarov, M., Taylor, J., Flibotte, S., LaBella, M., Pozniakovsky, A., Moerman, D.G., Jorgensen, E.M., 2014. Random and targeted transgene insertion in Caenorhabditis elegans using a modified Mos1 transposon. Nature Methods 11, 529–534.

Gauthier, K., Rocheleau, C.E., 2017. C. elegans Vulva Induction: An In Vivo Model to Study Epidermal Growth Factor Receptor Signaling and Trafficking., in: Wang, Z. (Ed.), ErbB Receptor Signaling. Methods in Molecular Biology. Humana Press, New York, NY.

Giordano-Santini, R., Milstein, S., Svrzikapa, N., Tu, D., Johnsen, R., Baillie, D., Vidal, M., Dupuy, D., 2010. An antibiotic selection marker for nematode transgenesis. Nature Methods 7, 721–723.

Hitoshi, N., Ken-ichi, Y., Jun-ichi, M., 1991. Efficient selection for high-expression transfectants with a novel eukaryotic vector. Gene 108, 193–199.

Hunt-Newbury, R., Viveiros, R., Johnsen, R., Mah, A., Anastas, D., Fang, L., Halfnight, E., Lee, D., Lin, J., Lorch, A., McKay, S., Okada, H.M., Pan, J., Schulz, A.K., Tu, D., Wong, K., Zhao, Z., Alexeyenko, A., Burglin, T., Sonnhammer, E., Schnabel, R., Jones, S.J., Marra, M.A., Baillie, D.L., Moerman, D.G., 2007. High-Throughput In Vivo Analysis of Gene Expression in Caenorhabditis elegans. PLOS Biology 5, e237.

Kage-Nakadai, E., Imae, R., Suehiro, Y., Yoshina, S., Hori, S., Mitani, S., 2014. A Conditional Knockout Toolkit for Caenorhabditis elegans Based on the Cre/loxP Recombination. PLOS ONE 9, e114680.

Kimble, J., Hirsh, D., 1979. The postembryonic cell lineages of the hermaphrodite and male gonads in Caenorhabditis elegans. Developmental biology 70, 396–417.

Kroetz, M.B., Zarkower, D., 2015. Cell-Specific mRNA Profiling of the Caenorhabditis elegans Somatic Gonadal Precursor Cells Identifies Suites of Sex-Biased and Gonad-Enriched Transcripts. G3: Genes|Genomes|Genetics 5, 2831–2841.

Lakso, M., Sauer, B., Mosinger, B., Lee, E.J., Manning, R.W., Yu, S.H., Mulder, K.L., Westphal, H., 1992. Targeted oncogene activation by site-specific recombination in transgenic mice. Proceedings of the National Academy of Sciences 89, 6232–6236.

Linkert, M., Rueden, C.T., Allan, C., Burel, J.-M., Moore, W., Patterson, A., Loranger, B., Moore, J., Neves, C., MacDonald, D., Tarkowska, A., Sticco, C., Hill, E., Rossner, M., Eliceiri, K.W., Swedlow, J.R., 2010. Metadata matters: access to image data in the real world. Journal of Cell Biology 189, 777–782.

Liu, K., Jin, H., Zhou, B., 2020. Genetic lineage tracing with multiple DNA recombinases: A user’s guide for conducting more precise cell fate mapping studies. Journal of Biological Chemistry 295, 6413–6424.

Luo, K.L., Underwood, R.S., Greenwald, I., 2020. Positive autoregulation of *lag-1* in response to LIN-12 activation in cell fate decisions during *C. elegans* reproductive system development. Development (Cambridge, England) 147, dev193482.

Madisen, L., Zwingman, T.A., Sunkin, S.M., Oh, S.W., Zariwala, H.A., Gu, H., Ng, L.L., Palmiter, R.D., Hawrylycz, M.J., Jones, A.R., Lein, E.S., Zeng, H., 2010. A robust and high-throughput Cre reporting and characterization system for the whole mouse brain. Nature Neuroscience 13, 133–140.

Maxwell, I.H., Harrison, G.S., Wood, W.M., Maxwell, F., 1989. A DNA cassette containing a trimerized SV40 polyadenylation signal which efficiently blocks spurious plasmid-initiated transcription. BioTechniques 7, 276–280.

Merritt, C., Rasoloson, D., Ko, D., Seydoux, G., 2008. 3’ UTRs Are the Primary Regulators of Gene Expression in the C. elegans Germline. Current Biology 18, 1476–1482.

Nagy, A., 2000. Cre recombinase: The universal reagent for genome tailoring. genesis 26, 99–109.

Qadota, H., Inoue, M., Hikita, T., Köppen, M., Hardin, J.D., Amano, M., Moerman, D.G., Kaibuchi, K., 2007. Establishment of a tissue-specific RNAi system in C. elegans. Gene 400, 166–173.

Ruijtenberg, S., van den Heuvel, S., 2015. G1/S Inhibitors and the SWI/SNF Complex Control Cell-Cycle Exit during Muscle Differentiation. Cell 162, 300–313.

Schindelin, J., Arganda-Carreras, I., Frise, E., Kaynig, V., Longair, M., Pietzsch, T., Preibisch, S., Rueden, C., Saalfeld, S., Schmid, B., Tinevez, J.-Y., White, D.J., Hartenstein, V., Eliceiri, K., Tomancak, P., Cardona, A., 2012. Fiji: an open-source platform for biological-image analysis. Nature Methods 9, 676–682.

Shin, H., Reiner, D.J., 2018. The Signaling Network Controlling C. elegans Vulval Cell Fate Patterning. Journal of Developmental Biology 6, 30.

Sternberg, P.W., 2005. Vulval development. WormBook : the online review of C. elegans biology, 1–28.

Tan, P.B., Lackner, M.R., Kim, S.K., 1998. MAP kinase signaling specificity mediated by the LIN-1 Ets/LIN-31 WH transcription factor complex during C. elegans vulval induction. Cell 93, 569–580.

Tenen, C.C., Greenwald, I., 2019. Cell Non-autonomous Function of daf-18/PTEN in the Somatic Gonad Coordinates Somatic Gonad and Germline Development in C. elegans Dauer Larvae. Current biology : CB 29, 1064–1072.e1068.

Tourasse, N.J., Millet, J.R.M., Dupuy, D., 2017. Quantitative RNA-seq meta-analysis of alternative exon usage in C. elegans. Genome Research 27, 2120–2128.

van der Vaart, A., Godfrey, M., Portegijs, V., van den Heuvel, S., 2020. Dose-dependent functions of SWI/SNF BAF in permitting and inhibiting cell proliferation in vivo. Science advances 6, eaay3823.

Wang, S., Tang, N.H., Lara-Gonzalez, P., Zhao, Z., Cheerambathur, D.K., Prevo, B., Chisholm, A.D., Desai, A., Oegema, K., 2017. A toolkit for GFP-mediated tissue-specific protein degradation in *C. elegans*. Development (Cambridge, England) 144, 2694–2701.

Ward, A., Liu, J., Feng, Z., Xu, X.Z.S., 2008. Light-sensitive neurons and channels mediate phototaxis in C. elegans. Nature Neuroscience 11, 916–922.

Watts, J.S., Harrison, H.F., Omi, S., Guenthers, Q., Dalelio, J., Pujol, N., Watts, J.L., 2020. New Strains for Tissue-Specific RNAi Studies in *Caenorhabditis elegans*. G3: Genes|Genomes|Genetics 10, 4167–4176.

Winston, W.M., Molodowitch, C., Hunter, C.P., 2002. Systemic RNAi in *C. elegans* Requires the Putative Transmembrane Protein SID-1. Science (New York, N.Y.) 295, 2456–2459.

Zhang, L., Ward, J.D., Cheng, Z., Dernburg, A.F., 2015. The auxin-inducible degradation (AID) system enables versatile conditional protein depletion in *C. elegans*. Development (Cambridge, England) 142, 4374–4384.

